# RhoG and Cdc42 can contribute to Rac-dependent lamellipodia formation through WAVE Regulatory Complex-binding

**DOI:** 10.1101/603340

**Authors:** Matthias Schaks, Hermann Döring, Frieda Kage, Anika Steffen, Thomas Klünemann, Wulf Blankenfeldt, Theresia Stradal, Klemens Rottner

## Abstract

Cell migration frequently involves the formation of lamellipodial protrusions, the initiation of which requires Rac GTPases signalling to heteropentameric WAVE regulatory complex (WRC). While Rac-related RhoG and Cdc42 can potently stimulate lamellipodium formation, so far presumed to occur by upstream signalling to Rac activation, we show here that the latter can be bypassed by RhoG and Cdc42 given that WRC has been artificially activated. This evidence arises from generation of B16-F1 cells simultaneously lacking both Rac GTPases and WRC, followed by reconstitution of lamellipodia formation with specific Rho-GTPase and differentially active WRC variant combinations. We conclude that formation of canonical lamellipodia requires WRC activation through Rac, but can possibly be tuned, in addition, by WRC interactions with RhoG and Cdc42.

## INTRODUCTION

Cell migration is essential for many physiological and pathological processes, such as embryonic development, immunity and metastasis [1–3]. Protrusion of the plasma membrane to enable cell migration is commonly achieved by the formation of thin branched arrays of actin filaments, called lamellipodia [4,5]. Small GTPases of the Rac family (i.e. Rac1, Rac2, Rac3 in mammals) are essential for lamellipodia initiation and maintenance, at least in part, by activation and continuous interaction through two binding sites on Sra-1 (or its orthologue PIR121) embedded into heteropentameric WAVE regulatory complex (WRC) [6,7]. The closest relatives of Rac GTPases are RhoG and Cdc42. Both fail to initiate lamellipodia in the absence of Rac expression [6], but can clearly promote Rac-dependent lamellipodia formation. This activity is thought to derive from crosstalk involving distinct Rac-GEF complexes [8–11]. The inability of RhoG and Cdc42 to initiate lamellipodia has hitherto been thought to be due to a lack of sufficient interaction with WRC. This is because both RhoG and Cdc42 show at best a weak interaction with Sra-1/PIR121 [6,12], and Cdc42, as opposed to Rac1, fails to activate native WRC *in vitro* [13].

Aside from Sra-1 (or PIR121), WRC is composed of four additional subunits – WAVE2 (or its paralogues WAVE1/WAVE3), the Sra-1/PIR121 interactor Nap1 (or Hem1 in the hematopoietic system), Abi1 (or Abi2/Abi3) and HSPC300 [14–17]. The Sra-1 subunit is “transinhibiting” the Arp2/3 complex-activating, so called WCA domain located on the C-terminal end of WAVE proteins, and Rac binding to Sra-1 outcompetes this inhibitory interaction to release the WCA domain, making it accessible for actin and Arp2/3 complex binding [18]. We have recently shown that the two aforementioned Rac binding sites on Sra-1/PIR121 [18,19] are essential for allosteric activation of WRC in vivo [7]. However, in spite of the previously proposed safe box model requiring two keys to allow for WRC activation to occur [19], we have found surprisingly specific physiological functions for the two sites in live cells. Whereas the low affinity A site is crucial for activation *in vivo*, the high affinity D site is contributing to the efficiency of lamellipodial protrusion, but by no means as important for WRC activation as the A site. Aside from the apparent critical function of Rac in WRC activation, it is less well established ifor if so to what extent Rac-WRC interactions also drive WRC recruitment to and accumulation in the lamellipodium. For instance, we have also found that lamellipodia formation can be initiated, in principle, without direct WRC-Rac interactions once WRC is rendered active, assuming at least that introduced, respective mutations of both A and D sites into active WRC abolished them entirely [7,19]. This is consistent with the fact that deleting the CAAX-box in Rac1, which is crucial for plasma membrane association, does impair, but not abolish lamellipodia formation in Rac1 KO fibroblasts [6]. In spite of the absence of an unequivocal, alternative mechanism of WRC recruitment, these data suggest that activation and lamellipodial targeting of WRC might potentially be separable. An example of such a separation clearly constitutes the related GTPase Cdc42, which mediates activation of its downstream effectors FMNL2 and −3 as prerequisite of their lamellipodial targeting [20,21], which however can fully occur upon removal of any potential GTPase interaction surface ([22] and unpublished data). Therefore, effector recruitment to lamellipodia is possible, in principle, without engagement of a given GTPase in spite of its established relevance in effector activation. Here, we have developed novel cell lines to compare the capability of lamellipodia formation by activated WRC in the absence or presence of endogenous Rac GTPases.

## RESULTS

### Rac – WRC interactions at the plasma membrane are dispensable for lamellipodia formation

We sought to test if plasma membrane insertion of the prenyl group of Rac1 is required for lamellipodia induction in B16-F1 cells. For this, we expressed constitutively active (Q61L), myc-tagged Rac1 or an identical construct lacking the CAAX-box essential for C-terminal prenylation in B16-F1 cells lacking Rac1/2/3 (clone#1; [7]). In analogy to our previously published experiments employing fibroblasts genetically deleted for Rac1 [6], deletion of the CAAX-box reduced but did not abolish lamellipodia formation in these conditions. More specifically, these structures were induced with overall slightly reduced frequency, and the majority of them (roughly 60%) appeared to be immature, as opposed to the majority of cells expressing full length, constitutively active (Q61L) Rac1 harbouring fully developed lamellipodia (Figures S1A and S1B). These data suggest that the same effects seen in Rac1^−/−^ fibroblasts [6] were not cell-type specific, and could not potentially be explained by remnants of Rac2 or −3 protein expressed perhaps at undetectable levels from respective genes not targeted in these fibroblasts.

In analogy, WRCs assembled in Sra-1/PIR121 KO cells (clone #3) harbouring a Sra-1 variant mediating constitutive WRC activation, but lacking functional Rac binding sites ([7] and Figure S1C) can still rescue lamellipodia formation, albeit at strongly compromised frequency [7]. Once formed though, and although compromised, these lamellipodia can still accumulate WRC at their tips (Figure S1D, see also [7]), and mediate continuous protrusion that is less smooth though than with lamellipodia driven by WRCs harbouring WT-Sra1 (Figure S1E). Together, both datasets thus suggest that although helpful, continuous Rac-WRC interactions at the plasma are not necessarily obligatory for lamellipodium protrusion.

### Generation of a cell line allowing further dissection of the Rac-WRC signalling module

Next we asked whether Rac is essential for WRC-mediated lamellipodia formation solely because of its essential function in WRC activation or because of serving additional functions. To test this, we had to develop cell systems in which endogenous WRC or Rac proteins could be replaced by active variants of each or functional deficiency mutants in a combinatorial fashion. To this end, we used CRISPR/Cas9 technology to disrupt Sra-1/PIR121 (KO clone #3) and Rac1, −2 and −3 (KO clone #1) encoding genes in B16-F1 cells individually [7] and in combination, giving rise to a novel cell line disrupted for all five genes (KO #3/11; Figure 1A). The latter now allows deciphering Rac/WRC signalling in more detail. In these cells lamellipodia formation is strikingly dependent on exogenous expression of Sra-1 and Rac1. While neither expression of EGFP as control, EGFP-Sra-1 nor myc-Rac1L61 alone facilitated lamellipodia formation in these cells, co-transfection of EGFP-Sra-1 and myc-Rac1L61 potently restored lamellipodia in these cells (Figures 1B and 1C). Notably, Rac1L61 expression in these WRC-deficient cells caused prominent plasma membrane blebbing (Figure 1B for representative image), reminiscent of our previous observations upon Rac microinjection upon WRC subunit knockdown [15]. This phenotype was robust and occurred at high frequency (data not shown), although a precise, mechanistic understanding of the phenomenon is currently lacking. Surprisingly, however, expression of EGFP-tagged Sra-1 WCA*, which should give rise to constitutively active WRCs, thus lacking the need for Rac-mediated WRC activation was incapable of driving lamellipodia formation in this cell line and conditions. If comparing this result with constitutively active WRCs compromised in Rac binding, but giving rise to inefficient lamellipodia formation in Sra-1/PIR121-KO cells ([7] and Figures S1C and S1D), two theoretical explanations for this discrepancy are thinkable: Firstly, the A+D site-mutated, active WRC used in Figure S1 can still inefficiently bind to endogenous Rac proteins present in these cells or secondly, Rac possesses additional, essential functions absent in cells harbouring active WRC but lacking endogenous Rac proteins (Figures 1B and 1C). Future experiments will have to distinguish between these possibilities.

**Figure 1.**
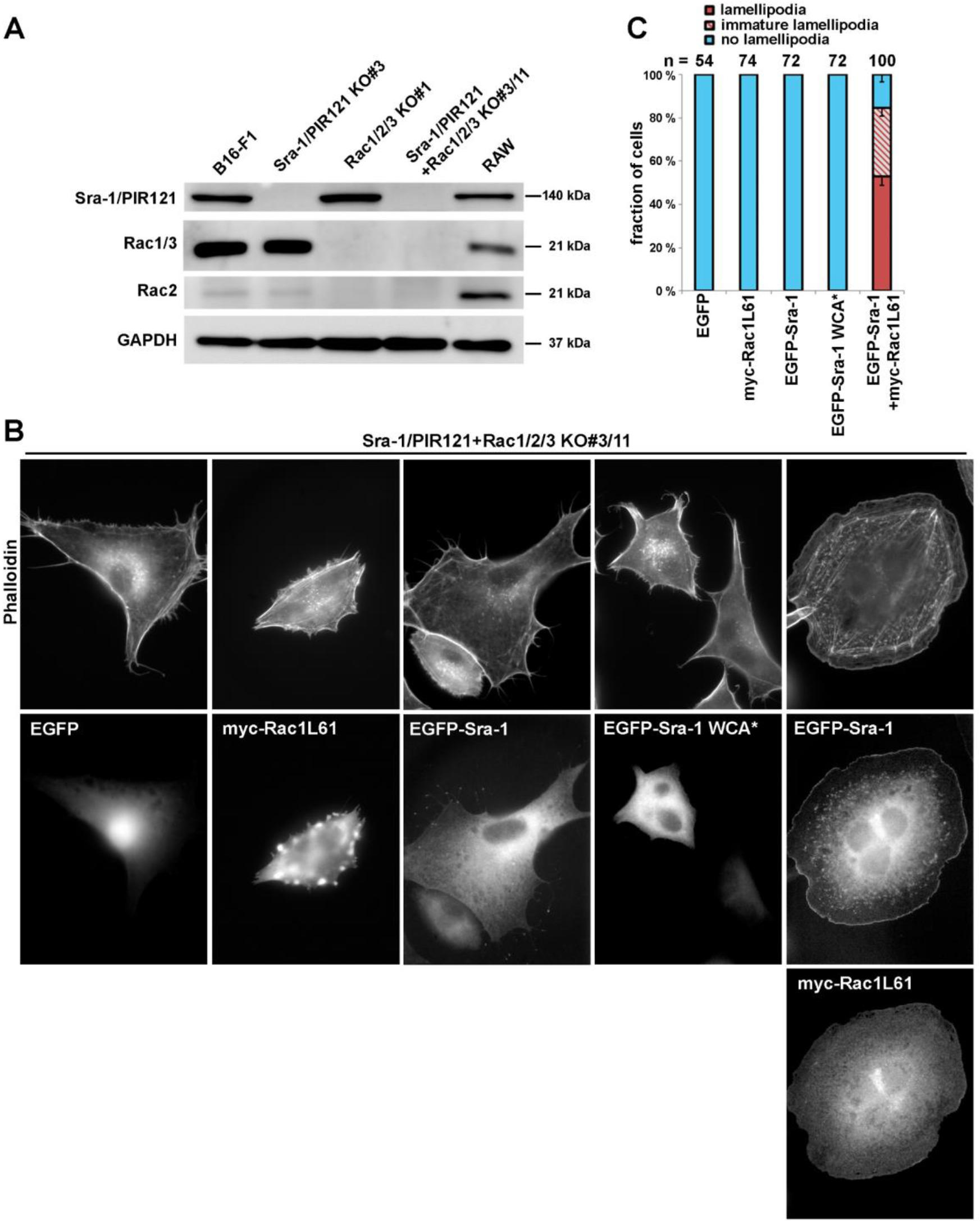
Generation of Sra-1/PIR121+Rac1/2/3 KO cells. (A) Cell lysates of indicated cell lines subjected to western blotting to probe expression levels of Sra-1/PIR121 and Rac GTPases. (B) Cell morphologies of Sra-1/PIR121+Rac1/2/3 KO cells (clone #3/11) expressing constructs as indicated. Panels in top row show stainings of the actin cytoskeleton with phalloidin, and middle and/or bottom row images show fluorescence of the same cells derived from either EGFP or anti-myc antibody stainings, as indicated. (C) Quantification of lamellipodial phenotypes. EGFP-Sra-1 WCA* denotes a construct rendering WRC constitutively active due to mutations relieving the autoinhibitory interaction of Sra-1 with the C-terminal WCA-domain (hence WCA* or active WCA) of WAVE [7,18]. Lamellipodial actin networks that were generally small, narrow, irregular or displayed multiple ruffles were defined as “immature lamellipodia”, as opposed to regular lamellipodia.

### Rac-related Rho GTPases fail to activate WRC, but can substitute for Rac in the presence of activated WRC

Up to this date, the literature contains conflicting results concerning the relevance of the closest relatives of Rac GTPases in mammals, in particular RhoG, but to a certain extent also Cdc42. In spite of prominent studies establishing functions for RhoG in signalling complexes operating upstream of Rac [10–12], RhoG has also already been concluded to contribute to fibroblast migration independent of Rac activation [23], although it has remained unclear how that occur mechanistically [24]. Of note, and again in full accordance with our previously published fibroblast data [6], our Rac1/2/3-deficient B16-F1 melanoma line previously failed entirely to form lamellipodia even upon expression of constitutively active RhoG (Figure 2A and B). Identical results were obtained with overexpressed, constitutively active Cdc42 but not Rac1, which robustly restored lamellipodia formation (Figures 2A and 2B), as expected [7]. Aside from the incapability of Cdc42 to induce lamellipodia, we found a prominent induction of stress fibres in these conditions (Figure 2A), which was not seen with RhoG. Such an activity is traditionally still mostly attributed to RhoA/B/C activity in the literature, and thus not followed up further in the context of this study. However, since our previous efforts allowed us to experimentally separate Rac-mediated WRC activation from other potential functions ([7] and see above), we wondered whether we might – in analogy to Rac - be able to establish connections between RhoG or Cdc42 activities and WRC function independent of Rac-mediated WRC activation. For this, we explored RhoG- or Cdc42-driven actin cytoskeleton remodelling in cells exclusively harbouring active WRC but lacking endogenous Rac GTPases (see Figure 1). And indeed, co-transfection of active (Q61L), myc-tagged RhoG or Cdc42 together with Sra-1 WCA* led to partial rescue of lamellipodia formation in Sra-1/PIR121+Rac1/2/3 KO cells (clone #3/11) (Figures 3A and 3C for quantitations). While RhoG was able to induce immature lamellipodia in more than 40 % of transfected cells, Cdc42 caused lamellipodia formation only in a small subfraction of cells (see Figures 3A, asterisks, and 3C for quantification). Nevertheless, both RhoG and Cdc42 caused accumulation of Sra-1 WCA* at sites of lamellipodia protrusion (Figure 3B). In contrast, replacing these GTPases by myc-tagged, constitutively active RhoA (G14V) failed to induce lamellipodia (Figure 3A, and C for quantification) and accumulation of EGFP-tagged Sra-1 WCA* at the cell periphery (Figure 3B), in spite of the GTPase clearly being active, as indicated by the expected simulation of stress fibres (Figure 3A).

**Figure 2.**
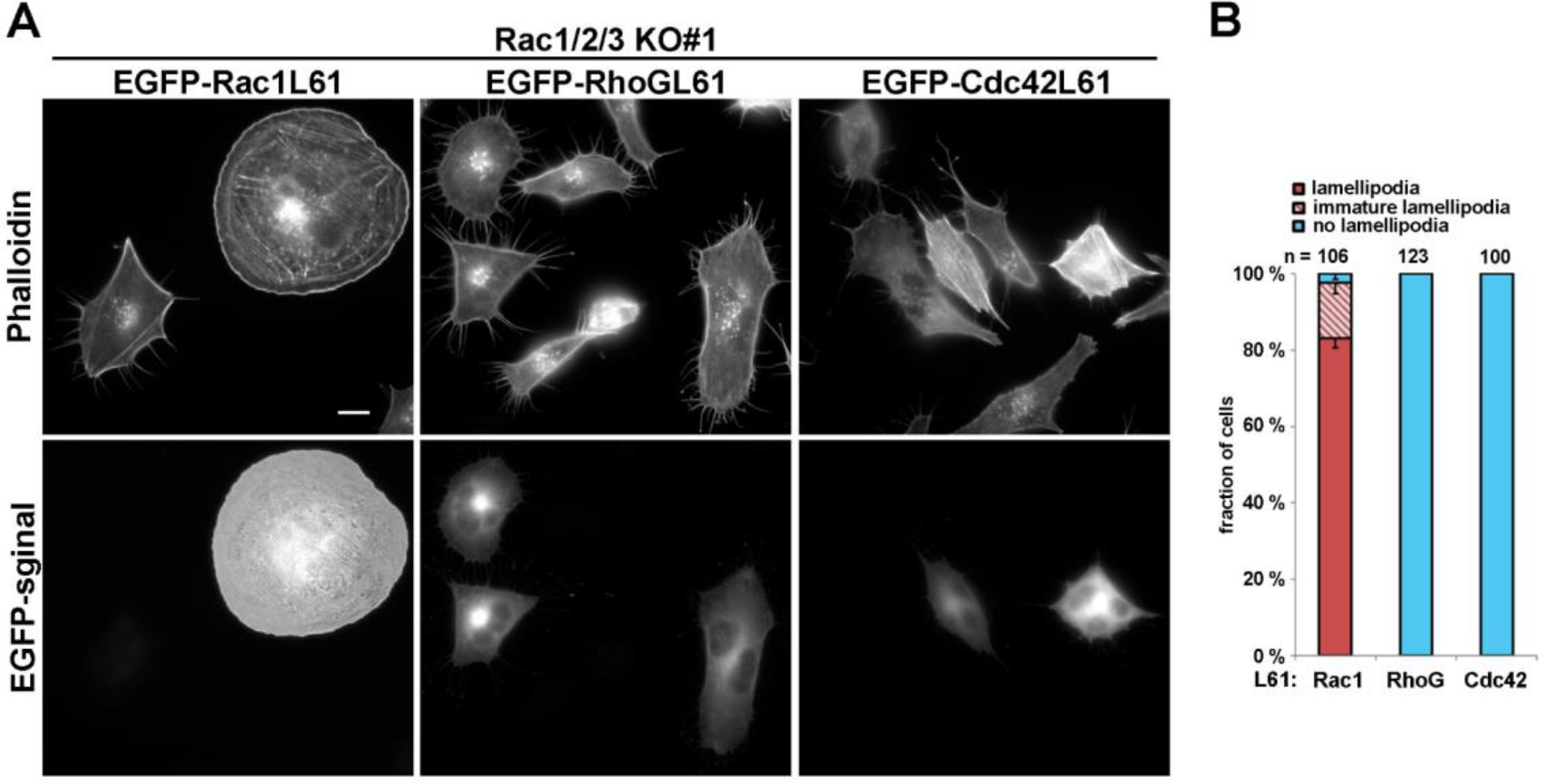
Rac-related GTPases RhoG and Cdc42 fail to induce lamellipodia in the absence of Rac expression. (A) Cell morphologies of Rac1/2/3 KO cells (clone #1) expressing EGFP-tagged GTPases as indicated. (B) Quantification of lamellipodial phenotypes was performed as described for Figure 1C.

**Figure 3.**
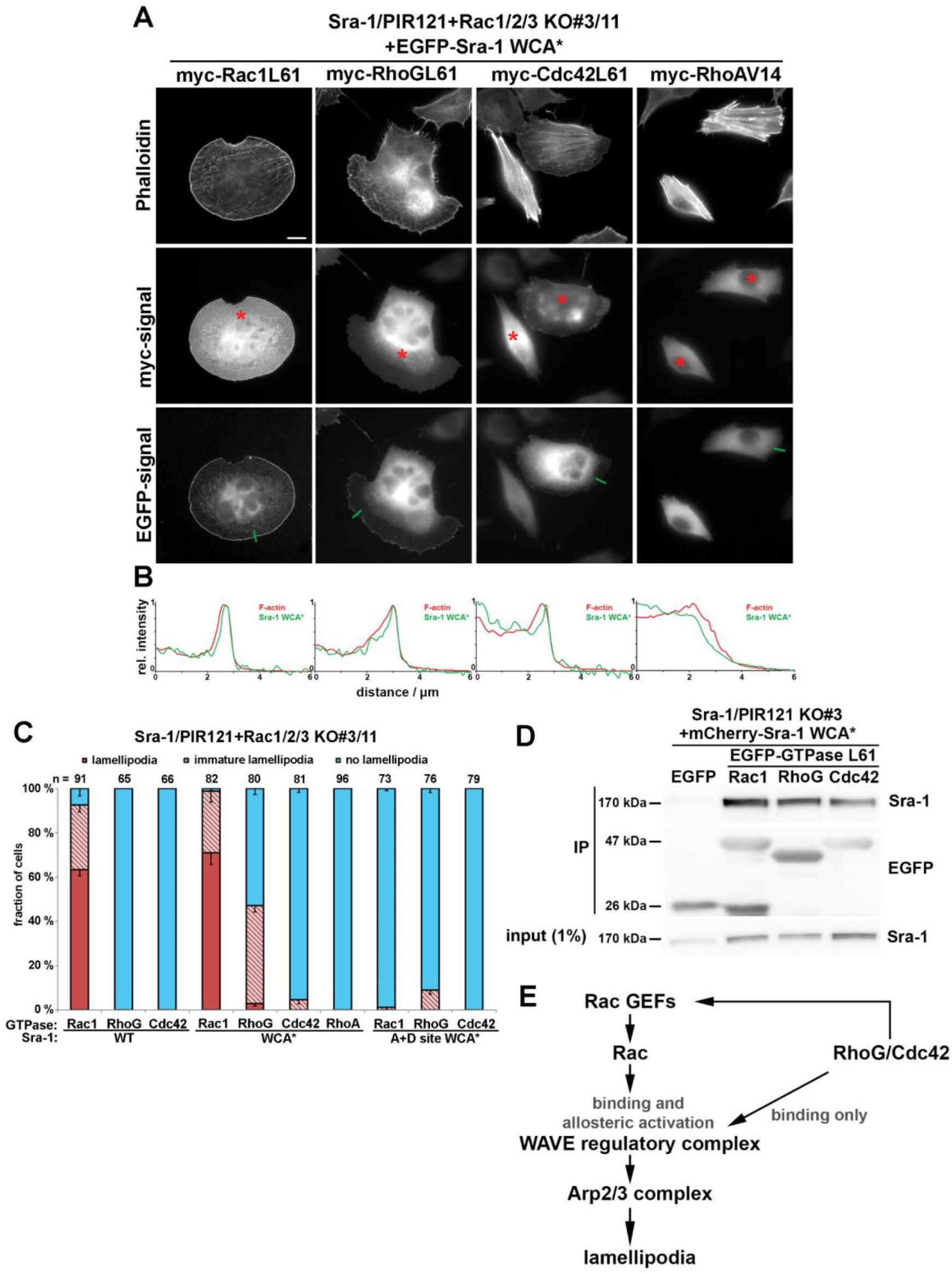
Induction of lamellipodia by RhoG or Cdc42 in conjunction with activated WRC, but in the absence of Rac expression. (A) Representative images of Sra-1/PIR121+Rac1/2/3 KO cells (clone #3/11) expressing EGFP-Sra-1 WCA* together with myc-tagged, small GTPases as indicated. Transfected cells identified by myc-staining (asterisks) were analysed concerning their cell morphologies (top row, actin filament staining with phalloidin) and the capability to accumulate active WRC (Sra-1 WCA*) at peripheral lamellipodia (B). Measurements shown in B were performed along line scans as shown in the images provided in A (green lines). (C) Quantification of lamellipodial phenotypes, done as described for Figure 1C. (D) Sra-1/PIR121 KO cells (clone 3) were co-transfected with mCherry-Sra-1 WCA* and either EGFP or constitutively active (Q61L), EGFP-tagged small GTPases, lysed and subjected to immunoprecipitation against EGFP. Note the complete absence of immunoprecipitation of Sra-1 upon co-expression of EGFP alone (negative control), and significant interactions of Sra-1 WCA* with Rac-1, RhoG and Cdc42. (E) Model for how RhoG and Cdc42 might regulate WRC and lamellipodia formation.

We confirmed that the ability of RhoG and Cdc42 to induce lamellipodia in this case was strictly dependent on constitutive WRC activity in this cell line, as both RhoG and Cdc42 failed to induce lamellipodia when co-expressed with WT-Sra-1 (Figure 3C). Finally, additional mutation of the two Rac binding sites on constitutively active WRC completely prevented (in case of Cdc42) or at least strongly inhibited (in case of RhoG) the induction of lamellipodia in these conditions (Figures 3A and 3C), strongly indicating that the action of RhoG/Cdc42 could mostly be explained by direct, (Rho-GTPase binding surface-dependent) interaction with WRC. In line with this, immunoprecipitation experiments clearly showed reasonably robust interactions of RhoG and Cdc42 with Sra-1 WCA*, albeit somewhat weaker perhaps than observed for Rac1 (Figure 3D).

All these results prompted us to compare the putative binding surfaces of RhoG and Cdc42 with the D site of Sra-1, as this site is the major affinity site for Rac GTPase [19]. Sequence alignments revealed that two (in case of RhoG) or three amino acids (in case of Cdc42) differed from Rac1 in the putative binding interface (Figure S2A). However, neither of these residues caused significant change in electrostatic surface potentials (Figure S2B). RhoA, on the contrary, showed obvious differences in the surface electrostatic potential in the putative binding interface, particularly caused by the glutamine to valine substitution at position 33 in RhoA, the analogous substitution of which in Rac1 (E31V) apparently interfered with proper lamellipodia formation [7,25]. Together, we can conclude that both RhoG and Cdc42 can specifically interact with WRC in a physiologically relevant manner once it has been activated by Rac1.

## DISCUSSION

Although puzzling initially, we have previously found and confirmed here using distinct experimental systems that Rac may not have to associate with the plasma membrane in order to activate or recruit WRC during lamellipodia initiation and maintenance. Yet, and not inconsistent with this view, Rac GTPases remain to be obligatory for both WRC activation and lamellipodia formation, as cells lacking both endogenous Rac GTPases and WRC can only form lamellipodia upon additional, specific manipulation. To our surprise, we establish for the first time here that lamellipodia formation is possible, in principle, without Rac GTPases, given that cells lack the need for WRC activation (because it is already activated or does not need to be activated), and that they over-express constitutively active variants of either RhoG or Cdc42. In other words, overexpression of RhoG, and to a lesser extent Cdc42 can cause the accumulation of constitutively active WRC even in the complete absence of endogenous Rac GTPases, presumably causing WRC-dependent Arp2/3 complex activation. This suggests that aside from previously established signalling crosstalk between Rac GTPases and RhoG/Cdc42 [8–10], the latter may directly contribute to the maintenance and/or activity of WRC at protruding lamellipodia edges once Rac has managed to activate individual WRCs.

Consistent with this view, we have previously found average turnover times for WRC subunits at the protruding plasma membrane that certainly fit the hypothesis of continuous activation events of Arp2/3 complexes, mediated by individual WRC units in a Rho GTPase binding-dependent fashion [7]. Slow turnover and molecular crowding of individual WRCs at the membrane likely contributes to efficient Arp2/3 activation at these sites (see also [26,27]), last, not least because efficient Arp2/3 complex activation was previously demonstrated to involve simultaneous engagement of two WCA domains [28–30]. Therefore, it seems plausible that efficient Arp2/3 complex activation at the lamellipodium tip coincides with WRC clustering, with the latter being affected, at least in part, by GTPase signalling.

The described, direct contribution of RhoG and Cdc42 to WRC-mediated actin remodelling is also found to occur in spite of at best weak or entirely non-specific interactions of RhoG and Cdc42 with wildtype WRC [6,12], as we show here that the situation changes dramatically if constitutively active WRC is used, which can bind quite efficiently to both Rac-related GTPases. This view is also confirmed by sequence alignments and structural considerations concerning effector interaction surfaces present on Rac1, RhoG and Cdc42 *versus* RhoA (Figure S2). Future structural studies will be needed to explain why the comparably robust interaction of RhoG and Cdc42 with Sra-1 observed here cannot occur with inactive WRC and/or translate into WRC activation.

Although the binding efficiency of RhoG and Cdc42 to active WRC appeared comparable in immunoprecipitation experiments, there was clearly measurable differences between the efficiency of the output response (lamellipodia in this case), the precise reasons for which remain to be determined. Yet, the vast majority of both RhoG and Cdc42-dependent lamellipodia formed in the absence of Rac GTPases were still immature, clearly illustrating relevance of Rac GTPases in formation and turnover of these structures beyond their established, essential function in WRC activation.

The novel cell system lacking endogenous Rac and WRC proteins (B16-F1 Sra-1/PIR121+Rac1/2/3 KO clone #3/11) also harbours the potential of emphasizing previously mentioned, but less well studied phenotypes caused by active, small GTPases, indicative for the commonly established, but chronically underestimated complexity arising from Rho-GTPase crosstalk [31,32]. For instance, without active WRC, Rac failed to induce lamellipodia, but caused prominent plasma membrane blebbing much less common to B16-F1 cells expressing endogenous WRC and thus capable of lamellipodia formation. Similar observations were previously reported for overexpression of the Rac effector loop mutant F37A [33], which we now consider to be impaired in driving lamellipodia formation as a result of compromised WRC interaction [7]. Whether this blebbing activity arises from intrinsic, Rac-specific features (i.e. requiring direct Rac-effector binding) or is a more indirect result of crosstalk to RhoA/B/C signalling remains unclear. Moreover, our preliminary observations also indicated that Cdc42 activities can funnel more prominently than usual into contractile stress fibre formation if Rac signalling (or lamellipodia formation) is missing. This could likely be mediated through signalling to MRCK kinases, previously shown to cooperate with Rho-ROCK signalling [34–36], which hitherto appeared less prevalent in our cell systems without compromised lamellipodia formation (see [37,38]). As the lamellipodia response seen with the Cdc42– (active) WRC combination was much less prominent than seen for the RhoG – (active) WRC couple, it is tempting to speculate that the stress fibre induction phenotype mentioned above might interfere with more robust lamellipodia formation. This might be explained perhaps by the widely accepted and long-standing antagonism between protrusion – (lamellipodia) *versus* contractility-dependent (stress fibres) processes [39,40]. Future experiments will have to reveal whether this assumption is correct.

Whatever the case, we are convinced that our approach of systematic generation and side-by-side comparison of Rho GTPase and effector knockouts in the same parental cell line will continue to unfold mechanistic insights into the intricacies of Rho signalling and crosstalk relevant for actin remodelling processes.

## MATERIALS AND METHODS

### Cell culture

B16-F1 cell line was purchased from ATCC (CRL-6323, sex:male). B16-F1 derived Sra-1/PIR121 KO cells (clone #3), as well as Rac1/2/3 KO cells (clone #1) were as described [7]. B16-F1 cells and derivatives were cultured in DMEM (4.5 g/l glucose; Invitrogen), supplemented with 10% FCS (Gibco), 2 mM glutamine (Thermo Fisher Scientific) and penicillin (50 Units/ml)/streptomycin (50 μg/ml) (Thermo Fisher Scientific). B16-F1 cells were routinely transfected in 35 mm dishes (Sarstedt), using 0.5 μg DNA in total and 1 μl JetPrime for controls, and 1 μg DNA in total and 2 μl JetPrime for B16-F1-derived knockout cells. After overnight transfection, cells were plated onto acid-washed, laminin-coated (25 μg/ml) coverslips and allowed to adhere for at least 5 hours prior to analysis.

### DNA constructs

pEGFP-C1 and –C2 vectors were purchased from Clontech Inc. (Mountain View, CA, USA). pEGFP-C2-Sra-1 was described previously [7] and corresponds to the splice variant *CYFIP1a*, sequence AJ567911. mCherry-tagged Sra-1 was generated by swapping EGFP with mCherry, kindly provided by Dr. Roger Tsien (University of California at San Diego, La Jolla, California, USA) using NheI/BsrGI restriction sites. pRK5-myc-Rac1L61 and pRK5-myc-RhoAV14 were kindly provided by Alan Hall and Laura Machesky (CRUK Beatson Institute, Glashow, UK). Cdc42L61 (placental isoform) was synthesized by Eurofins Genomics and cloned into pRK5-myc and pEGFP-C1 vectors. pEGFP-C1-Rac1L61 and pRK5-myc-RhoGL61 were as described [6,7]. For generation of pEGFP-C1-RhoGL61 the RhoGQ61L, the corresponding DNA fragment fragment immobilized from pRK5-myc-RhoGL61 with BamHI/EcoRI was ligated into pEGFP-C1 vector digested with BglII/EcoRI. pRK5-myc-Rac1L61-ΔCAAX was generated by site directed mutagenesis using 5’-GAGGAAGAGAAAATGACTGCTGTTGTAAGTC-3’ as forward primer. The fidelity of all constructs was verified by sequencing.

### CRISPR/Cas9-mediated genome editing

B16-F1 cells lacking functional *CYFIP1* and *CYFIP2* genes, as well as *Rac1*, *Rac2* and *Rac3* genes were generated by treating Sra-1/PIR121 KO cells (clone #3) with pSpCas9(BB)-2A-Puro (PX459) vectors targeting Rac1, Rac2 and Rac3 genes. Specifically, cells were co-transfected with plasmids targeting ATGCAGGCCATCAAGTGTG (Rac1/2) and ATGCAGGCCATCAAGTGCG (Rac3) genomic regions as described [7]. After puromycin selection of transfected cells (3 days), cells were extensively diluted and a few days later, macroscopically visible colonies picked, to obtain single cell-derived clones. Derived cell clones already lacking Sra-1/PIR121 were screened for the additional absence of Rac expression by Western Blotting (see Figure 1).

### Western blotting

Preparation of whole cell lysates was performed essentially as described [7]. Western blotting was carried out using standard techniques. Primary antibodies used were Sra-1/PIR121 [15], Rac1/3 (23A8, Merck), Rac2 [6], GAPDH (6C5, Calbiochem) and GFP (clones 7.1 and 13.1, Roche). HRP-conjugated secondary antibodies were purchased from Invitrogen. Chemiluminescence signals were obtained upon incubation with ECL™ Prime Western Blotting Detection Reagent (GE Healthcare), and were recorded with ECL Chemocam imager (Intas, Goettingen, Germany).

### Immunoprecipitation

For EGFP-immunoprecipitation experiments, Sra-1/PIR121 KO cells (clone #3) co-expressing EGFP alone or EGFP-tagged variants of constitutively active (Q61L) GTPases together with mCherry-tagged Sra-1 WCA* were lysed with lysis buffer (1% Triton X-100, 140 mM KCl, 50 mM Tris/HCl pH 7.4/ 50 mM NaF, 10 mM Na_4_P_2_O_7_, 2 mM MgCl_2_ and Complete Mini, EDTA-free protease inhibitor [Roche]). Lysates were cleared and incubated with GFP-Trap agarose beads for 60 min. Subsequently, beads were washed three times with lysis buffer lacking Triton X-100 and protease inhibitor, mixed with Laemmli buffer, boiled for 5 min and subjected to Western Blotting.

### Fluorescence microscopy, phalloidin and antibody stainings and quantification

B16-F1-derived cell lines were seeded onto laminin-coated (25 μg/ml), 15 mm-diameter glass coverslips and allowed to adhere for at least 5 hours. Cells were fixed with pre-warmed, 4% paraformaldehyde (PFA) in PBS for 20 min, and permeabilized with 0.05% Triton-X100 in PBS for 30 sec.

PFA-fixed cell samples following transfections with plasmids mediating expression of EGFP-tagged proteins were counterstained with ATTO-594-conjugated phalloidin (1:200). For stainings with myc antibodies, permeabilized cells were blocked with 5% horse serum and 1% BSA in PBS, followed by staining with monoclonal anti-myc antibody (9E10; undiluted, home-made hybridoma supernatant). Primary antibodies were visualized with Alexa Fluor 350-coupled anti-mouse IgG. Linescans were generated using MetaMorph software by drawing lines (width of 15 pixels) from inside the cell and across the lamellipodium.

### Time-lapse microscopy

Live cell imaging shown in Figure S1D was done with Sra-1/PIR121 KO #3 cells transfected with respective EGFP-tagged Sra-1 variants and migrating on laminin-coated glass coverslips (25 μg/ml). Cells were observed in μ-Slide 4 well (Ibidi), and maintained in microscopy medium (F12 HAM HEPES-buffered medium, Sigma), including 10% FCS, 2 mM L-glutamine and 1% penicillin/streptomycin. Conventional video microscopy was performed on an inverted microscope (Axiovert 100TV, Zeiss) equipped with an HXP 120 lamp for epifluorescence illumination, a halogen lamp for phase-contrast imaging, a Coolsnap-HQ2 camera (Photometrics) and electronic shutters driven by MetaMorph software (Molecular Devices). Live cell images were obtained using a 100 x/1.4 NA Plan apochromatic oil objective. Kymographs were generated using MetaMorph software by drawing lines from inside the cell and across the lamellipodium.

### In silico-comparison of binding interfaces of small GTPases to the D site of Sra-1

Sequence alignments were carried out using http://www.uniprot.org. The structure of RhoG was predicted with Phyre2 [41]. For comparison of surface electrostatic potentials of different small GTPases, indicated structures were superimposed with Rac1 occupying the D site of Sra-1 [7,19] using CCP4MG.

## Acknowledgements

This work was supported by grants within the framework of a graduate program supported by the Deutsche Forschungsgemeinschaft (DFG), called GRK2223/1 (to K.R. and W.B). We also thank Brigitte Denker for excellent technical assistance, Prof. Laura Machesky (CRUK Beatson Institute, Glasgow, UK) for Rho-GTPase expression constructs and Prof. Baoyu (Stone) Chen (Iowa State University, U.S.A.) for fruitful discussions.

## SUPPLEMENTARY FIGURES LEGENDS

**Figure S1.**
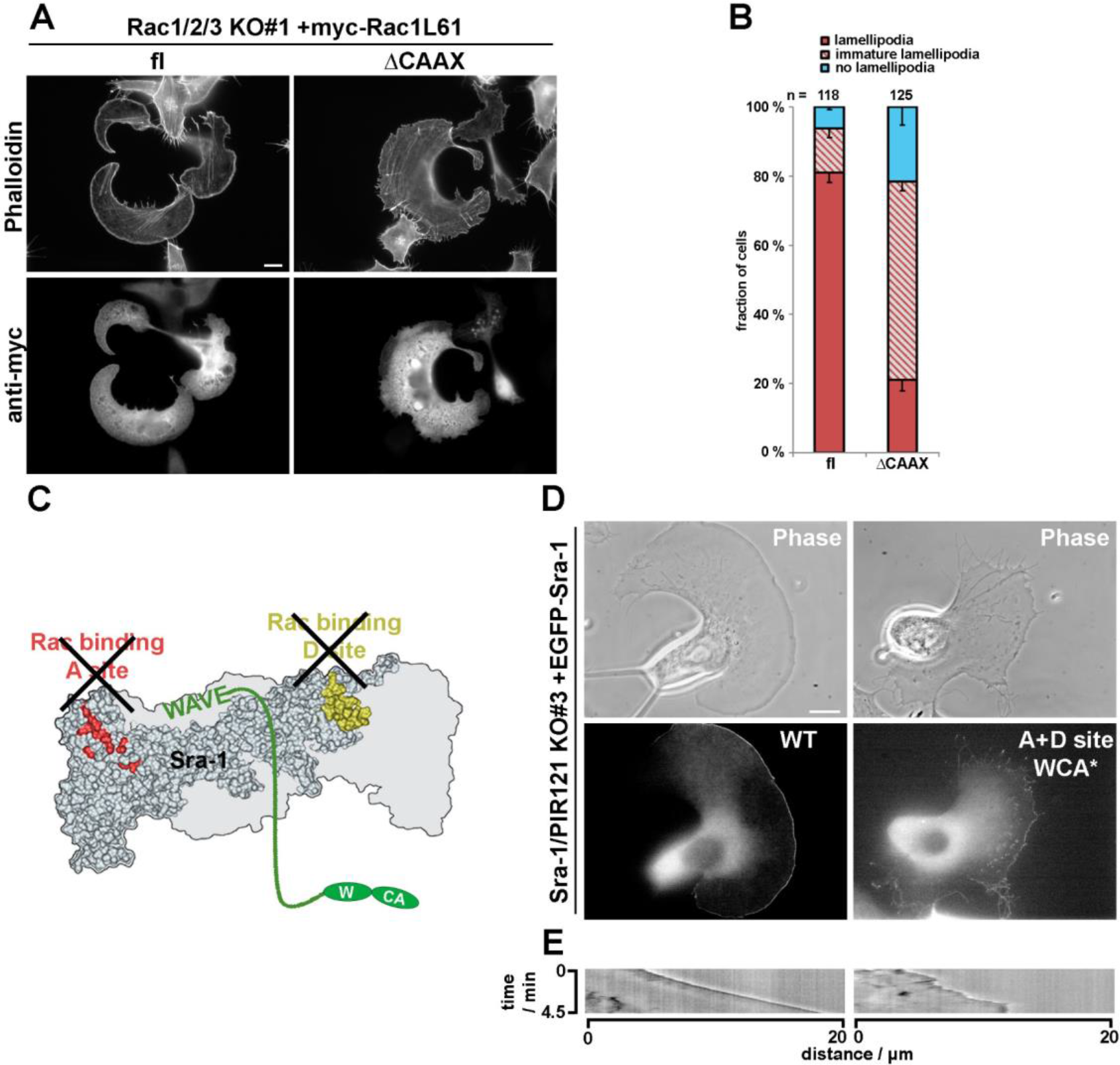
Contribution of the membrane-interacting region in Rac1 to lamellipodia formation. (A) Cell morphologies and lamellipodial phenotypes of Rac1/2/3 KO cells (clone#1) transfected with myc-tagged Rac1L61 variants and stained for myc and the actin cytoskeleton by phalloidin. (B) Quantification of lamellipodial phenotypes performed as described for Figure 1C. (C) Structural cartoon of the Sra-1 construct used in D and mediating assembly into a constitutively active WRC that lacks functional Rac binding sites (see also [7]). (D) Life cell imaging of Sra-1/PIR121 KO cells (clone #3) re-expressing indicated Sra-1 constructs. (E) Kymograph analysis showing membrane protrusion induced by respective construct.

**Figure S2.**
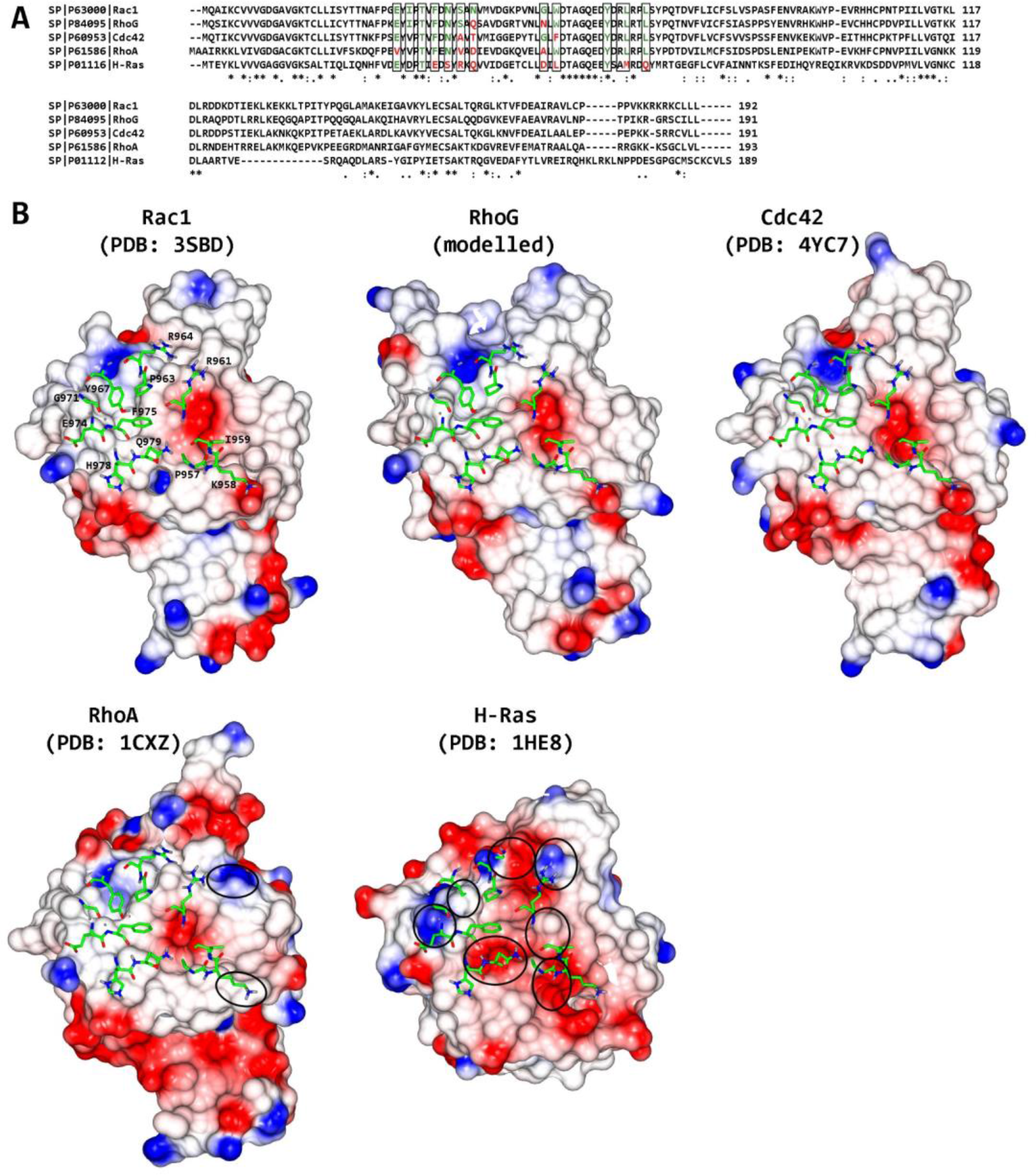
Comparison of binding interfaces of small GTPases to the D site of Sra-1. (A) Sequence alignment of human GTPases. Rectangles mark amino acids that contact the D site of Sra-1. Sequence identity to Rac1 is indicated by green, difference by red colouring of respective amino acids. (B) Surface electrostatic potential of small GTPases capable of contacting the D site of Sra-1. Structures of GTPases were superimposed with Rac1 occupying the D site. Key residues in Sra-1 contacting Rac1 are shown as sticks. Differences in the surface electrostatic potential in the putative binding interface or steric conflicts are encircled. Such differences seen in RhoA likely abolished potential responses comparable to RhoG or Cdc42 (see Figure 3). Note that the differences in Ras were so significant (7 circles in total) that we did not consider the latter to be of any relevance for lamellipodia formation *in vivo*.

